# Convergent Genomic Signatures of Selection in Thar: Genetic trade-offs for desert survival

**DOI:** 10.64898/2026.11.17.767204

**Authors:** Sachin M. Rathod, Rishabh Jain, Mitali Mukerji

**Author notes:** These authors contributed equally (Sachin M. Rathod, Rishabh Jain).

## Abstract

**Background:** Desert environments provide a setting for natural biological experiments to cope up with simultaneous stressors of extreme temperatures, aridity, UV exposure, and limited food resources. Biological adaptations that support living in such an environment could become liabilities when the environment and lifestyle change due to gene-environment mismatch. We explored whether genetically and culturally diverse endogamous native populations of the Thar exhibit convergent signatures of selection to desert conditions and whether the selected loci have implications in traits and disease susceptibilities relevant to non-native transitions.

**Results:** We report 348 high-confidence variants across 45 candidate genes from a study on 176 individuals from eight genetically distinct Thar populations. These genes identified from humans and other desert inhabitants display convergent evolution with multiple selection scans (PBSn1, Tajima’s D, iHS, and XPEHH) in diverse Thar populations. Prominent genes include *ZNF385D, SGCZ, SARDH, SLC12A3*, etc. Selected variants essential in the desert environment were associated with complex traits like lipid metabolism, hypertension, renal and cardiovascular diseases, and addiction in GWAS and PheWAS. Noteworthy, in a syntenic locus, eQTLs in *SARDH*, which modulates DBH implicated in addiction, suggest a potential link with survival advantage in an uncertain desert environment. Likewise, a selected variant (rs7204044) in *SLC12A3*, implicated in hypertension in non-desert populations, seems to be a key to water-salt homeostasis in desert natives.

**Conclusions:** These findings suggest that variants linked to adaptive traits in desert conditions could be linked to maladaptation in an altered environment. This has potential health consequences in climatic and lifestyle transitions among native inhabitants.

## Introduction

The desert environment imposes a combination of stressors, including extreme fluctuating temperatures, water scarcity, high solar radiation, and limited food availability, on its inhabitants. Despite challenges and adversities, diverse plants and animals have evolved physiological, behavioral, and molecular mechanisms that support survival under arid stress conditions (1–3). Genomic and transcriptomic studies of camels, desert rodents, foxes, and other desert animals have identified candidate genes and pathways associated with water and ion balance, energy metabolism, thermoregulation, oxidative stress, and cellular maintenance (4–6). The recurrence of related physiological pathways across evolutionarily distant species suggests that similar environmental pressures could work through shared biological processes, although the specific genes and variants involved may vary (7,8). These provide a foundational basis for investigating core genomic mechanisms that may contribute to human adaptation to desert environments.

Human populations have inhabited diverse ecological settings, and population-genomic studies have identified signatures of adaptation to high-altitude dietary transitions, infectious diseases, and other environmental pressures (9–11). However, genomic studies on human populations residing in hot deserts are relatively understudied compared to other extreme environments. A recent study on Turkana pastoralists in East Africa has provided evidence of candidate genomic signatures associated with life in an arid environment, including findings related to water stress (12). However, the genetic underpinnings of long-term human habitation in hot deserts remain incompletely understood, particularly in South Asia. An additional challenge is that demographic history, endogamy, genetic drift, and population structure generate patterns that resemble selection, identifying plausible adaptive candidates. Consequently, such studies need to be undertaken in genetically diverse populations residing in the same environment, along with complementary selection statistics and orthogonal functional evidence.

The Thar Desert in northwestern India provides an opportunity to investigate this question in a setting characterized by aridity, intense seasonal heat, high solar radiation, and water scarcity. The region supports diverse communities with distinct occupational practices and social histories, including pastoralists, artisans, semi-nomadic, and rural and urban populations. Endogamy, migration, and differences in livelihood and settlement practices may have contributed to genetic differentiation among these communities and disrupted their shared regional environment (13–16). A recent population genomic study by our group elucidated the genetic structure and the effects of migration, occupational identity, and demographic history on diverse Thar communities (17). Building on this population genetic context, the next question is whether the genomes of Thar communities also harbor signatures of selection in genes related to physiological processes relevant to desert living. Addressing this question requires distinguishing signals associated with population history from candidates that warrant further investigation for their possible functional relevance.

In this study, we investigated candidate genomic signatures of desert adaptation in Thar populations using a genome-wide approach to candidate genes, focusing on desert-related stress responses and physiological processes, informed by evidence from genomics, transcriptomics, and physiological studies on desert-adapted organisms. Genetic associations with complex traits were also explored for possible evolutionary trade-offs in a non-native environment. This study revealed evidence of convergent evolution across human populations with diverse ancestral histories, as well as between humans and other desert-dwelling species. We identified new loci that have not been previously implicated in human desert adaptation, a significant number of which were associated with complex diseases and traits. These findings provide a basis for future investigations into how adaptations to extreme environments can impact health outcomes amidst climate change and lifestyle transitions, particularly in the inhabitants of the Thar Desert.

## Materials and Methods

### Curation of the candidate genes reported for desert adaptation

A literature survey with keywords like desert adaptation, signature of selection, and pathways such as salt metabolism, renal function, skin physiology, and thermoregulation associated with desert stressors was conducted to curate genes from human and desert-dwelling animal studies (Figure 1). The desert inhabitants include rodents (6,8,18–25), camels (5,26–30), and other desert-adapted mammals such as fat-tailed sheep, other livestock, and North African foxes (4,31–35,35–40). Genes from human genomic studies of populations inhabiting hot and arid environments, including Turkana pastoralists (12), Australian Aboriginals, and other African populations, were also included (41–54) (Supplementary Table 1). We also curated genes from genomic, transcriptomic, and functional studies conducted in response to desert-associated stressors, i.e., heat, temperature fluctuations, osmotic stress, dehydration, and food deprivation. Genes were prioritized if they were i) reported across independent studies, ii) significantly expressed under desert stress conditions, and iii) functionally linked to pathways critical for desert survival. This reduces biases from a single study. This approach was preferred over a genome-wide scan to improve statistical power to detect the signal in biologically relevant genes and to explore convergence of adaptation among desert inhabitants.

**Figure 1.**
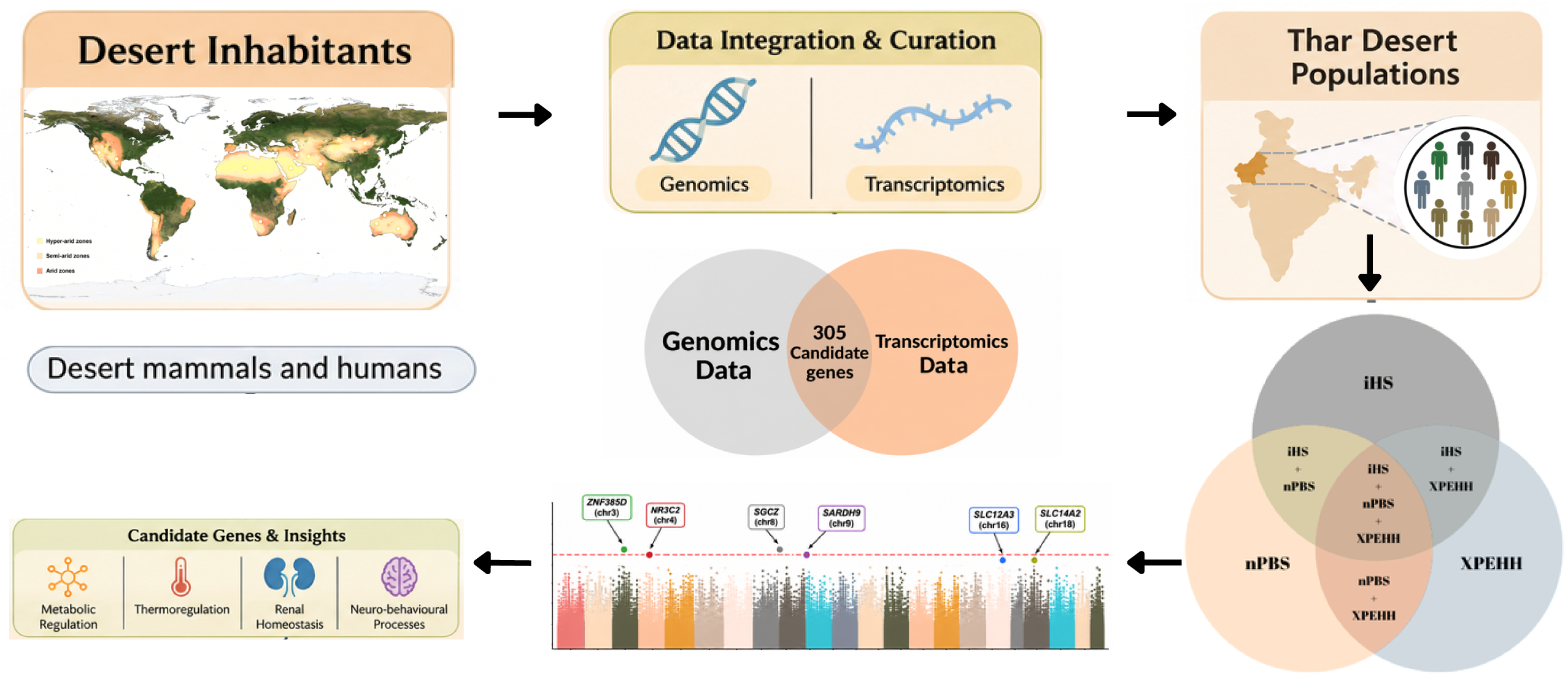
Schematic overview of the integrative approach for identifying candidate genes associated with desert adaptation through the identification of overlapping candidates across genomic and transcriptomic datasets and selection scans.

### Quality control, phasing, and imputation

We used the GenomeStudio v2 pre-QC’d genotype data from our previous study, which included 176 individuals and ∼730k markers across eight populations (17). Quality control was performed using PLINK1.9 (--hwe 1-e07, --geno 0.10, --mind 0.15) (55) to retain ∼498 markers. Furthermore, to improve genome-wide variant density and haplotype resolution beyond the array, we imputed our retained markers using the next-generation imputation platform Michigan Imputation Server Minimac4 (56). For quality control, phasing and imputation were performed internally with a reference panel from 1KGP South Asian (SAS) Phase 3 Version 5 (hg19), and an R2 filter of 0.3 was applied to retain well-imputed variants.

Downstream analysis of the imputed Thar genotype dataset was integrated with the filtered 1KGP hg19 WGS and HGDP datasets (57). Filtered and high-quality phased variants common to both datasets were extracted for intersecting markers using “*isec -n=2 -w1*- and merged using the “*merge -m none*” function implemented in BCFtools (58).

### Datasets

We analyzed genome-wide genotype data from 185 individuals belonging to nine endogamous communities representing artisan, pastoralist, semi-nomadic, and nomadic groups that were extensively characterized in our recent study of the genetic landscape of the Thar ecoregion (17). We integrated the Thar dataset with publicly available reference population datasets from the 1000 Genome Project (1KGP) (59) and the Human Genome Diversity Project (HGDP) (60,61) for comparative selection scan analysis.

### Multi-method selection scan for identification of signatures of selection

To identify the genetically proximate non-Thar South Asian reference population, we used data on pairwise F_ST_ and outgroup f3 analyses from our previous study (17). Pairwise F_ST_ estimates (62) give genetic differentiation between populations, and outgroup f3 (63,64) quantifies shared genetic drift between a target and reference population using a distantly related outgroup. Based on the concordance between pairwise F_ST_ and outgroup f3 estimates, we identified the genetically closest South Asian population for each Thar population group and used it as the comparative reference group in selection analyses.

We used complementary allele-frequency-based and haplotype-based selection-scan methods to detect the signature of selection in Thar populations. These include normalized population branch statistics (PBSn1) (44,65), Tajima’s D (66), integrated haplotype score (iHS) (53), and cross-population extended haplotype homozygosity (XP-EHH) (67). Selection scans were performed on 305 curated candidate genes in each Thar population against its genetically proximate South Asian reference populations, Gujarati (GIH), Punjabi (PJL), Pathan, and outgroup Yoruba (YRI), for lineage-specific analysis.

For PBSn1, pairwise F_ST_ values were calculated on the merged Thar-1KGP (87K markers) and Thar-HGDP (73K markers) datasets using PLINK v1.9 for Thar-reference, Thar-YRI, and reference-YRI comparisons. PBSn1 was computed as described in Yi et al. 2010, Malaspinas et al. 2016, and Veilleux et al. 2023 (44,65,68), with negative F_ST_ values set to zero before PBS calculation. Higher PBSn1 values indicated greater lineage-specific differentiation in the target Thar population. Tajima’s D was calculated using vcftools (69) with a 5 kb window for the 45 top genes with imputed SNPs. This was computed separately for each population with a selection threshold (Tajima’s D value below −2).

The haplotype-based analyses were performed in R with the rehh package using default parameters (70,71). iHS was calculated for each Thar population for SNPs with MAF >5%, which were treated as core SNPs. XP-EHH was calculated for each Thar population as the target, with GIH, PJL, and Pathan as the corresponding reference populations. Additionally, Extended Haplotype Homozygosity (EHH) decay was examined around the top candidate *ZNF385D* (rs1841916), *SLC14A2* (rs12455117), *THADA* (rs61483132), *TMTC1* (rs1438515), and *SLC12A3* (rs7204044) from the stringent selection to assess the extent and consistency of the haplotype-based signals across the Thar and reference populations.

To identify robust selection candidates, the results from the PBSn1, iHS, and XP-EHH were integrated. Variants occurring within the top 1% of at least two selections and recurring across at least two comparative populations were prioritized. These criteria were applied to reduce the reliance on any single statistic and to identify candidate variants supported by multiple independent tests (72). Only variants meeting these criteria were carried forward for functional annotation and possible biological interpretation.

### Annotation and Characterization of Candidate Variants

To assess the biological relevance of the top 348 high-confidence variants identified from the integrated selection scan, functional annotation, pathway enrichment, and disease and trait associations were carried out using existing genomic resources and databases. Variants were annotated using SNPnexus v4 (73) and the Ensembl Variant Effect Predictor (VEP) (74) to determine the functional consequences, regulatory status, and function category. Further, genes associated with the top variants were analyzed for functional enrichment using the ToppFun module of the TopGene Suite (75) to assess the gene-level function, Gene Ontology, and relevance to desert adaptation.

The phenotypic association summary statistics for the 348 high-confidence selected variants were investigated using publicly available data. PheWas was performed using the *phewas()* function in the ieugwasr R package (76,77) against the IEU OpenGWAS database (https://opengwas.io/), querying five data batches: ebi-a (NHGRI-EBI GWAS Catalog) (78), eqtl-a (eQTL Catalogue) (79), ieu-b (IEU curated multi-trait collection), ukb-b (UK Biobank Neale Lab Round 2), and ukb-e (Pan-UK Biobank ancestry-stratified GWAS) (80), included for its non-European strata relevant to South Asian allele frequencies. Queries were run in chunks with error handling, and associations were retained at a threshold of p<1×10, allowing for both suggestive and established associations.

To complement this with higher-confidence evidence, the NHGRI-EBI GWAS Catalogue was separately queried using gwasrapidd (81). Additionally, a subset of the variants were cross-checked in GWASatlas for phenotypic and pleiotropic associations (82). The subset of variants from the eQTL catalogue of OpenGWAS was cross-validated against tissue-specific cis-eQTL data from the GTEx portal (83). For convergent selection and Phewas/GWAS colocalization, regional linkage disequilibrium (LD) was mapped using LDBlockShow v1.41 for top genes (84).

## Results

### Selection scan and identification of high-confidence convergent selection signals

By applying the top 1% threshold for PBSn1, iHS, and XP-EHH, followed by filtering signals in at least two selection scan methods, 348 high-confidence SNPs across eight Thar populations were identified (Figures 2-3), mapping to 45 protein-coding genes. 22 SNPs topped the list in all analyses (PBS+IHS+XPEHH) across the Thar population (Supplementary Table 2). The UpSet plot in Figure 2 represents the intersections of pairwise comparisons between Thar and three comparator populations. For example, 138 SNPs were common across selection scan comparisons with all reference populations. Notably, the largest number of SNPs were observed in *SGCZ* (n= 84) and ZNF385D (n= 74), with signals recurring across multiple populations and combinations of selection scans. Other notable genes showing recurrent signals included *CSMD1, GPC6, CACNA1E, CRADD, SARDH, BCL2, NR3C2, PPARGC1A, and TMTC1* (Supplementary Tables 3 and 4). The most prominent variants were rs1027345 from *ZNF385D*, rs1438151 from *TMTC1*, and rs4858395 from *ZNF385D*, which were featured across all sets.

**Figure 2.**
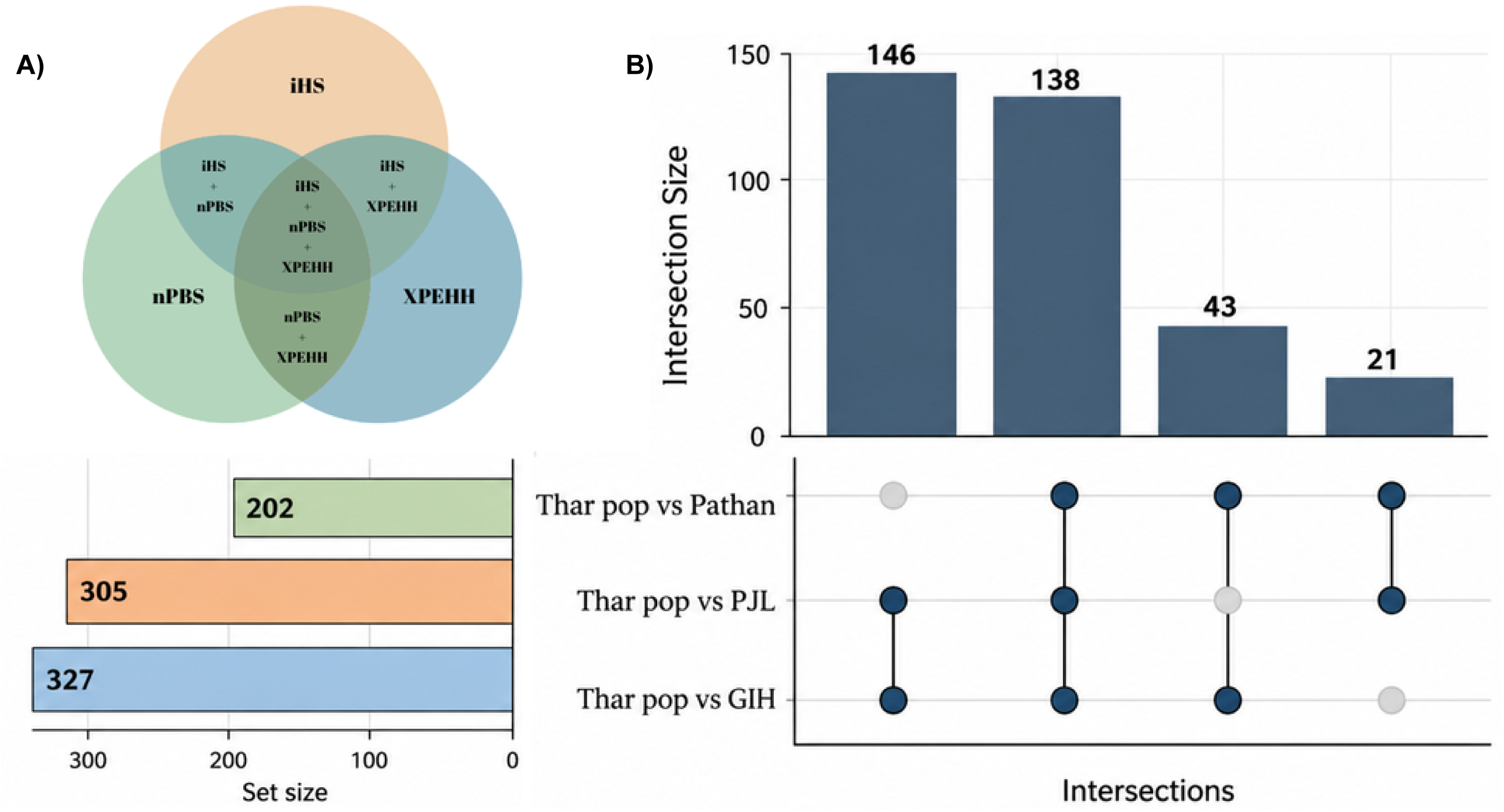
Identification of high-confidence candidate SNPs with selection criteria. A) Venn diagrams show SNP overlap among selection methods (PBSn1, iHS, and XP-EHH). B) UpSet plots illustrate their recurrence across comparative population analyses (GIH, PJL, and Pathan). Only recurrent overlapping SNPs were retained as high-confidence candidates.

**Figure 3.**
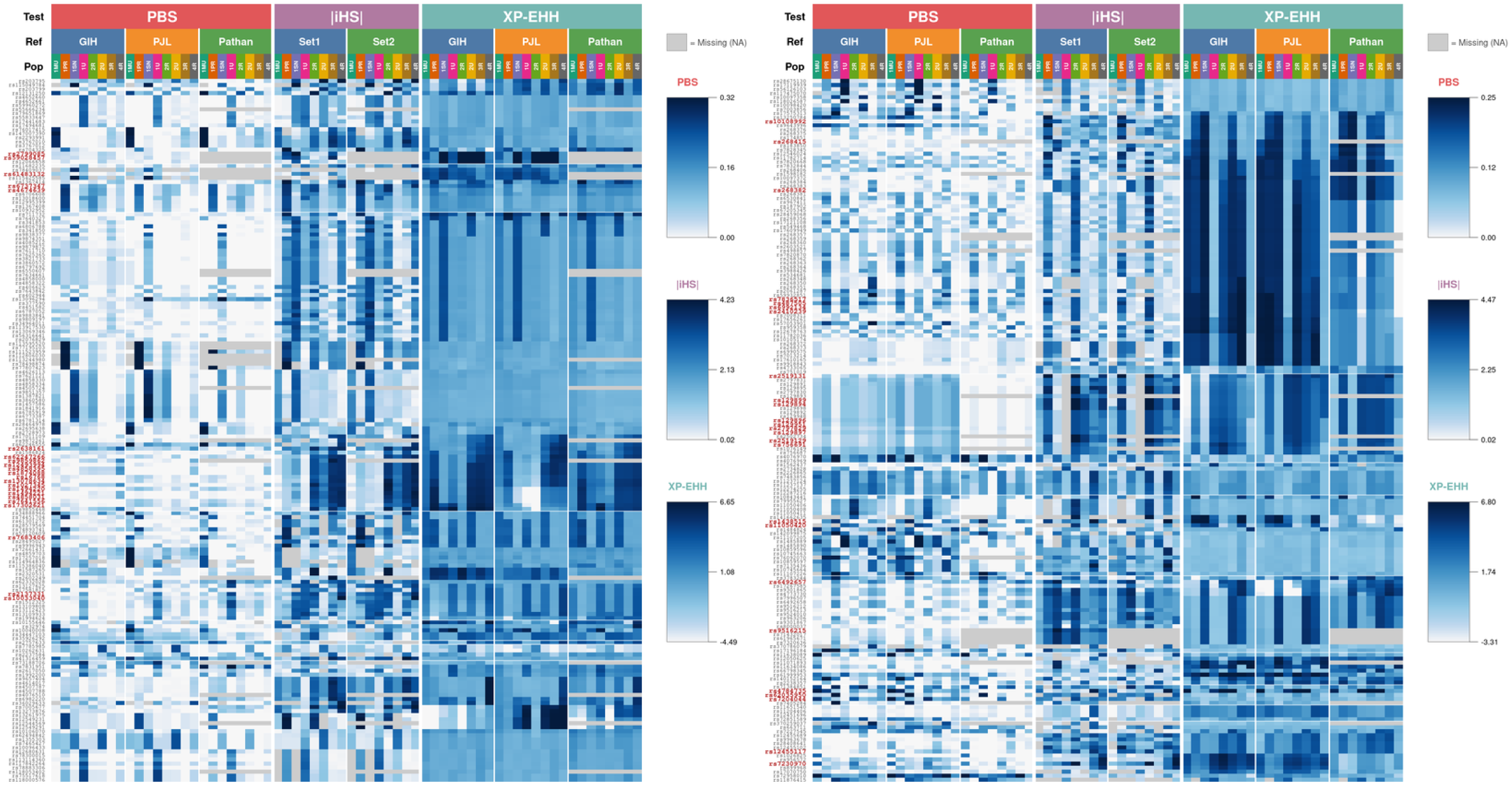
Heatmap of selection scores for the top 348 candidate SNPs, three selection tests (PBSn1, iHS, and XP-EHH) for each Thar population and their respective comparative populations. A color gradient ranging from white to blue represents low to high selection scores, respectively, for each test. SNPs highlighted in red denote variants recurrently identified under selection in more than two Thar populations.

To explore the potential influence of demographic history on patterns of genetic variation, Tajima’s D was calculated across the genomic regions of the top 45 candidate genes that deviate from the neutral expectations. Signatures of selection (<-2) were identified in 28 genes which overlap with most of the genes described above and a few other genes like *TBXAS1*, *SLC12A1*, and *NOTCH1*, etc. (Supplementary Table 5).

The allele-specific EHH decay profile provides additional insight into the haplotype structure surrounding the candidate variants across Thar and reference populations (Supplementary Figure 1). EHH decay profile around the derived allele of rs12455117 (*SLC14A2*) showed comparatively slow decay in the haplotype in the Thar population compared to reference populations. Overall, the EHH profiles provide additional support for extended haplotypes around the derived alleles at several candidate loci, although the strength of this pattern differs across populations.

### Functional annotation and biological classification of the candidate genes

Functional annotation of the 348 high-confidence SNPs identified in 45 protein-coding genes showed that most variants were located in intronic regions. (n=323) Overall, the variants were predominantly located in non-coding regions, with limited representation in protein-coding sequences. (Supplementary Figure 2, Supplementary Table 6).

Functional enrichment analysis using ToppFun with Q-value Bonferroni < 0.05 revealed significant ontologies with lipid measurements, waist-hip ratio, tissue morphogenesis, chemical and cellular homeostasis, and transmembrane transport. Enrichment of mouse phenotypes related to distal tubule morphology and renal transport was evident in kidney function (Supplementary Figure 3). Based on the function and their physiological relevance, we hypothesized that the selected genes could be broadly grouped into six major biological axes: endurance and resilience, cellular stress tolerance, metabolic conservation and adaptive thermogenesis, water-salt homeostasis, skin barrier and extracellular matrix organization, and neuro-behavioral functions (Figure 4).

**Figure 4.**
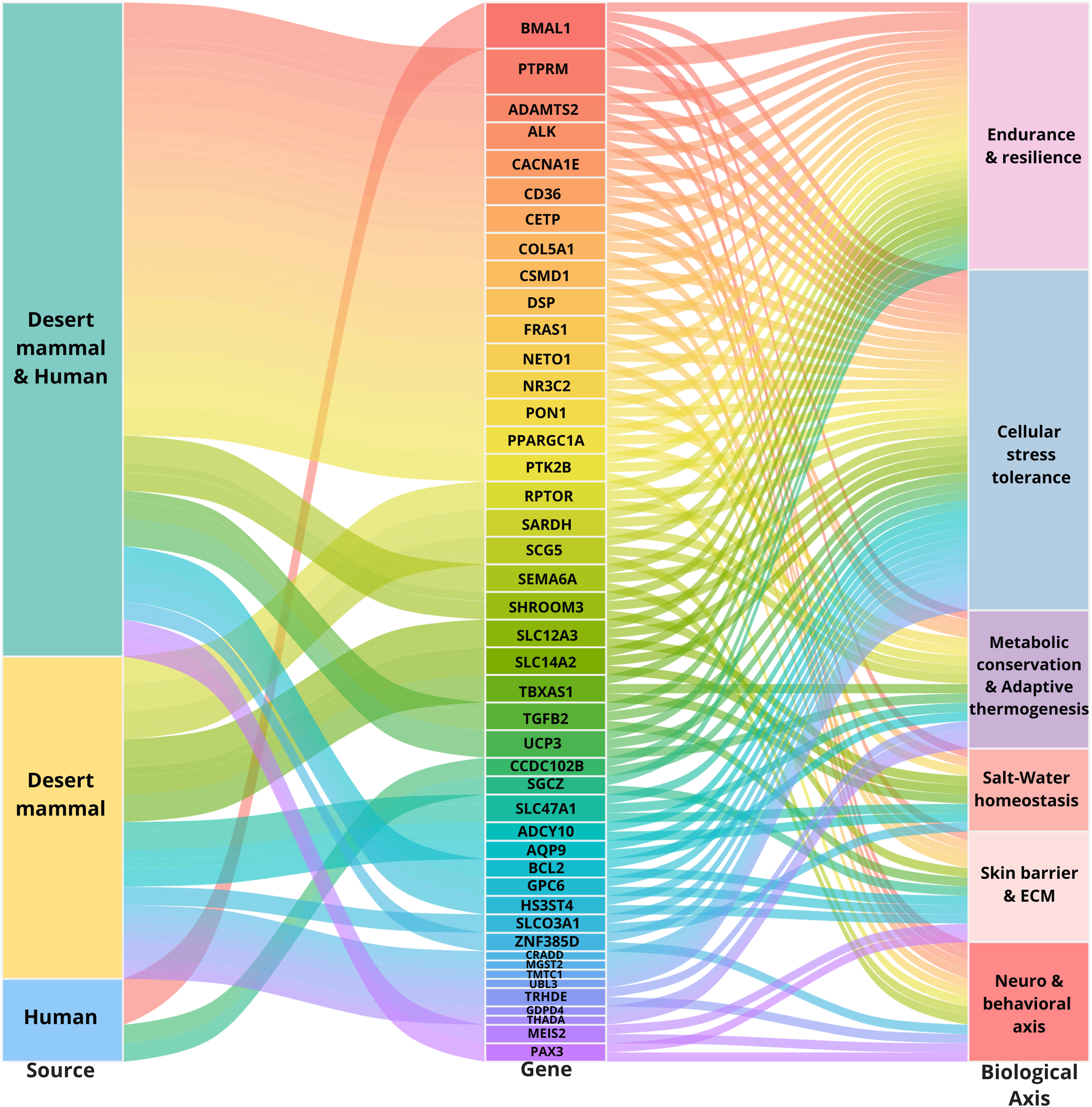
Distribution of the 45 candidate genes identified through selection-scan analyses. Genes are linked to supporting literature from desert mammals and humans and are mapped to predicted biological axes associated with desert adaptation.

Furthermore, PheWAS analysis was conducted using ieugwasr and gwasrapidd, supplemented by the NHGRI-EBI GWAS catalogue, and we identified phenotypic associations for selected candidate SNPs. The *ZNF385D* variant rs1874088 was associated with longevity and appeared across seven population-comparison combinations. The *SGCZ* variant rs7003856 was associated with neurofibrillary tangles, whereas *GPC6* rs6492657 was associated with heel bone mineral density. Among the candidate genes implicated in renal transport and homeostasis, *SLC14A2* rs12455117 was found to be associated with hypertension. Additionally, the variants rs62035950, rs4784735, and rs7204044 of *SLC12A3* were found to be associated with HDL cholesterol (Supplementary Figure 4, Supplementary Table 7,8).

The 19 out of 20 high-confidence SNPs identified in *SARDH* were reported as cis-quantitative trait loci (cis-QTLs) for the adjacent genes *DBH* and *DBH-AS1*, with the strongest associations in subcutaneous adipose, visceral omental adipose, and skin tissues (Supplementary Figure 5). The variants rs2519131 and rs129899 showed consistently negative normalized effect sizes (NES) for both genes, indicating associations with reduced expression. These results suggest that the associated alleles were linked to reduced expression of both genes in the queried tissues. Additionally, *SLC12A3* (rs7204044) showed a strong association with HDL cholesterol levels in GWASatlas, with a p<1×10 ^31^ (Supplementary Figure 6).

On examining the regional LD-based analysis, a distinct haplotype architecture was revealed. The *SARDH* showed a compact 6 kb high LD block around rs2519131, while *SLC12A3* contained two distinct blocks, including rs7204044. *SGCZ* exhibited a fragmented 609 kb structure containing six major LD blocks with 84 candidate SNPs, including rs7003856. *ZNF385D* spanned 887 with three major LD blocks, one containing candidate variant rs1874088 (Supplementary Figure 7)

## Discussion

This study builds on our earlier reported genetically distinct native communities of the Thar Desert, who have inhabited this environmentally challenging region for thousands of years. Our study reveals signatures of selection in pathways involving 45 coding genes. Many of these are implicated in desert adaptation in other mammals but remain unexplored in human populations (Table 1). Several variants under selection in the desert populations are also associated with complex diseases and traits in non-native populations. This exploratory, candidate gene-based study offers insights into potential genetic trade-offs in environments where populations have transitioned.

**Table 1.** Summary of the top 45 candidate genes reported in human populations and desert-adapted animal species. Genes sharing the same sources are grouped in one row. Human populations: SAF, Southern Africa/Kalahari (Ju|’hoansi, Xoo, Herero, Khoe-San); EAF, East Africa (Mursi, Turkana, Amhara); SAH, Sahel/North Africa/Sahara (Fulani, North African Berbers); PAF, Pan-African (Bantu-speaking populations); SAS, South Asia (Indian populations, OGWIP from the IGV dataset); OCE, Oceania (Aboriginal Australians); SIB, Siberia (Naukan Yup’ik, Maritime Chukchee); 1KG, 1000 Genomes Project panel; ANC, ancient human genomes. Desert animals: ROD, desert rodents (cactus mouse, northern three-toed jerboa, Siberian jerboa, midday jird, Roborovski dwarf hamster, banner-tailed kangaroo rat, desert-adapted house mouse, olive grass mouse, canyon mouse, deer mouse); CAM, camelids (Bactrian camel, dromedary); LIV, livestock (goat breeds, Barki goat, Barki sheep, Sheep HapMap breeds, Mongolian cattle); POU, poultry (Saudi Arabian chicken); ODM, other desert mammals (North African foxes, Tarim red deer). Study type: G, genomics; T, transcriptomics; G+T, genomics and transcriptomics.

| **Gene** | **Human populations** | **Desert animals** | **Study** | **Functional association of selected variant (PheWAS / GWAS)** |
|---|---|---|---|---|
| **Genes reported in both human populations and desert animals** |  |  |  |  |
| ***CACNA1E*** | SAF, EAF, SAH | ROD | G+T | cis-eQTL for CACNA1E expression |
| ***TGFB2*** | SAF, EAF, SAH, SAS, OCE | ROD | G+T | — |
| ***ALK, GPC6, MEIS2*** | ANC | ROD | G+T | GPC6: bone mineral density |
| ***PAX3*** | SAH, SAS | ROD | G+T | Hair color (pigmentation) |
| ***ZNF385D*** | EAF, ANC | ROD | G+T | Longevity |
| ***PPARGC1A*** | EAF, OCE, SIB, 1KG | ROD, LIV | G+T | cis-eQTL for PPARGC1A expression |
| ***SHROOM3, FRAS1, ADAMTS2*** | SAS | ROD | G+T | SHROOM3: kidney function (cystatin C), gut microbiome<br>ADAMTS2: height, body fat percentage |
| ***NR3C2*** | SAF, EAF, SAH | ROD, CAM | G+T | Pulse rate |
| ***SEMA6A*** | EAF | ROD | G+T | Corpus callosum volume; SEMA6A/COMMD10 eQTL |
| ***DSP*** | SAS, ANC | ROD | G+T | Lung function (FEV1/FVC) |
| ***CD36*** | EAF, PAF, OCE, SIB, 1KG | ROD | G+T | — |
| ***PON1, UCP3, CETP*** | EAF, OCE, SIB, 1KG | ROD | G+T | UCP3: mean platelet volume; UCP3/UCP2 and neighboring-gene eQTL |
| ***CSMD1*** | EAF, SAH, ANC | ROD, WLD | G+T | Age at menarche, height |
| ***PTK2B*** | SAF, EAF, SAH, SAS, OCE, SIB, 1KG | ROD | G+T | Eosinophil count, BMI; TRIM35/EPHX2 eQTL |
| ***COL5A1, BCL2*** | SAF, EAF, SAH, SAS | ROD | G+T | BCL2: serum protein levels, bone mineral density |
| ***HS3ST4*** | SAF, SAH, ANC | ROD | G+T | — |
| ***PTPRM*** | SAF, EAF, SAH, ANC | ROD, POU | G+T | — |
| ***NETO1*** | OCE | ROD | G+T | — |
| **Genes reported in human populations only** |  |  |  |  |
| ***SGCZ*** | EAF | — | G | Neurofibrillary tangles |
| ***BMAL1*** | SAF, EAF, SAH | — | G | ARNTL eQTL; gut microbiome |
| ***CCDC102B*** | EAF, SAH | — | G | TMX3 eQTL |
| **Genes reported in desert animals only** |  |  |  |  |
| ***ADCY10, THADA, TMTC1, UBL3, SCG5, SLCO3A1, SLC12A3, SLC47A1, RPTOR, SLC14A2*** | — | ROD | T | ADCY10: neutrophil and white blood cell counts; MPZL1 eQTL<br>THADA: blood pressure/hypertension, HbA1c, blood cell counts, age at menarche<br>TMTC1: cis-eQTL for TMTC1 expression<br>UBL3: SLC7A1 eQTL<br>SCG5: ARHGAP11A/GOLGA8N eQTL<br>SLCO3A1: kidney function (eGFR reduction); SLCO3A1 eQTL<br>SLC12A3: lipid metabolism (HDL cholesterol, apolipoprotein A-I, lipoprotein particles)<br>SLC47A1: kidney function (creatinine, eGFR); eQTL<br>RPTOR: adiposity (BMI, weight, fat mass)<br>SLC14A2: myopia, breast cancer, bone mineral density |
| ***MGST2*** | — | CAM | T | — |
| ***TBXAS1*** | — | ROD, POU | G+T | — |
| ***SARDH*** | — | ROD, CAM | T | DBH/DBH-AS1 eQTL |
| ***GDPD4*** | — | ROD, CAM, WLD | G+T | PAK1/INTS4 eQTL |
| ***TRHDE*** | — | ROD, LIV | G+T | — |
| ***CRADD*** | — | ROD, LIV, WLD | G+T | Height, sitting height, lean mass, physical function; SOCS2 eQTL |
| ***AQP9*** | — | ROD | G+T | Lipid metabolism (triglycerides in HDL, apolipoprotein A) |

Selection scans using complementary methods across candidate genomic regions identified 348 high-confidence SNPs across 45 protein-coding genes in Thar populations, compared with genetically and geographically proximal populations from Non-Thar regions. Functional enrichment analyses grouped these genes to six interconnected biological axes involving water-salt homeostasis, metabolic regulation and thermogenesis, cellular stress responses, endurance and resilience, skin physiology, and neurobehavioral functions (Figure 4). These biological processes may collectively contribute to survival under desert conditions. The enrichment of genes involved in renal and electrolyte transport is particularly relevant in the context of prolonged heat exposure and limited water availability. Similarly, genes associated with energy metabolism, thermoregulation, and cellular maintenance may contribute to maintaining physiological homeostasis under environmental stress. Notably, the predominance of non-coding variants among the identified selection signals suggests potential modulation of gene expression rather than changes in protein sequence. Furthermore, the reported association of some variants with multiple phenotypes in the PheWAS database suggests potential pleiotropic effects and highlights the possibility of genetic trade-offs associated with adaptation. Together, these findings suggest that coping with diverse environmental stressors in the desert may involve coordinated changes across multiple physiological systems rather than isolated adaptive responses.

A substantial proportion of candidate genes showing selection signals have been reported in non-human studies, although some candidates have also been reported in other desert dwellers. For instance, *SGCZ* and *ZNF385D*, with 84 and 74 candidate SNPs, have also been reported in East African Turkana pastoralists, where *SGCZ* encodes zeta-sarcoglycan, a transmembrane component of the sarcoglycan. It is involved in the maintenance of the extracellular matrix and contributes to muscle-cell membrane stability (85). Whereas, ZNF385D encodes a zinc-finger protein with a potential role in DNA binding and transcriptional regulation. Recent evidence suggests its role in silencing insulin secretion in a beta-cell model (86). This overlap raises the possibility of convergent genomic responses to similar environmental pressures across geographically distinct desert-adapted populations (12). Noteworthy, 20 variants spanning a 6 kb region from another candidate gene, *SARDH*, involved in one-carbon metabolism reported in desert mammals, showed selection signals across some genetically diverse Thar populations (1PR, 2R, 4R, and 2U). Most of these variants were non-overlapping, with only rs2519131 shared across all four. This pattern may indicate a combination of shared and population-specific selective responses, although the functional significance of these variants remains to be established. Another notable candidate was *THADA*, which was prioritized and selected based on its elevated expression across a multi-tissue transcriptome of desert-adapted cactus mice in water-deprived states (20). Interestingly, *THADA* has also been widely reported in the adaptive response to extreme cold conditions through non-shivering adaptive thermogenesis in humans (87,88). This gene has been additionally associated with hypertension and type 2 diabetes in GWAS (89,90). These observations highlight the potential relevance of *THADA* to metabolic and thermoregulatory processes across contrasting environmental conditions, while also raising the possibility of a broader role in coping with diurnal temperature fluctuations characteristic of desert populations.

Additional, selection was also observed in the genes sodium transporter (*SLC12A3*), urea transporter (*SLC14A2*), and mineralocorticoid receptor (*NR3C2*). These are biologically plausible candidates for desert adaptation given their involvement in salt water homeostasis, fluid balance, and renal physiology (91–93). Besides desert humans, these genes are also reported in cactus mice, kangaroo rats, and camels, suggesting potentially converging pathways across desert-dwelling species. (5,6,19–21). The *SLC12A3* variant (rs7204044) is reported in Phewas as associated with HDL and cholesterol and also for hypertension in East Asian populations (94–96). This selected variant may play an important role in desert inhabitants to maintain electrolyte balance and blood pressure under conditions of prolonged water loss. Its association raises the possibility of a link with lipid metabolism. Together, these findings highlight *SARDH* and *SLC12A3* as promising candidates connecting regulatory variation, renal physiology, and metabolic processes with potential desert adaptation.

One of the most striking observations was the signature of selection upstream of the Sarcosine dehydrogenase (*SARDH*) locus, adjacent to the Dopamine B hydroxylase (*DBH*) locus and its antisense transcript (*DBH-AS1*) (Supplementary Figure 8). *SARDH* is a key mitochondrial enzyme that converts sarcosine to glycine, which links glycine betaine, which plays a key central role in one carbon metabolism. Levels of *SARDH* could modulate intracellular osmolyte homeostasis during cycles of severe dehydration and rapid rehydration in the desert environment (7,97).

All 19 out of 20 variants mapping to the 6-kb region at the *SARDH* locus are cis-eQTLs for *DBH* and *DBH-AS1* in adipose (subcutaneous and visceral) as well as sun-exposed and unexposed skin. The derived alleles observed in the Thar populations were significantly associated with decreased expression. Although expression was highest in the liver, these variants were not identified as eQTLs in this tissue. *DBH* catalyzes the conversion of dopamine to norepinephrine and could thus influence catecholamine balance and sympathetic responses (98).

Reduced *DBH* levels have been associated with neuropsychiatric traits, addiction, pathological gambling, and reward-seeking behavior in studies of non-desert populations. Altered reward responses as well as reduced sympathetic activation could conceivably contribute to coping with the physiological and resource-related uncertainties in the desert environment. Interestingly, corroborating results were observed with prominent enrichment around amphetamine abuse and addiction in GO analysis. However, the effects of variants and physiological consequences remain to be established experimentally (98). Synteny of DBH and *SARDH* across other desert mammals suggests a conserved requirement of pathways associated with sympathetic regulation, thermoregulation, and osmoregulation in desert survival and subsistence.

This exploratory study highlights the possibility that genetic variations potentially advantageous under desert conditions may have different consequences under non-desert environments. Variations in metabolic and thermoregulatory pathways may interact with dietary changes, reduced physical activity, and changes in traditional occupation and urbanization that might be reflected in differences in adiposity and biochemical profiles across Thar communities and provide relevant context for examining and estimating these relationships. Though the current findings may not establish a direct association between identified selection and contemporary disease risk, further genotype-phenotype and gene-environment analyses are required to investigate these potential evolutionary trade-offs.

Overall, this study provides an integrated framework for investigating population structure and potential genomic signatures of desert adaptation in the Thar populations. The convergence of the selection statistics, haplotype structure, and functional annotation highlights several candidate genes and interconnected physiological pathways for further investigation and follow-up studies.

## Conclusion

This study identifies the candidate signature of selection across desert species and also across the genes involved in metabolic regulation, thermoregulation, renal homeostasis, and neuro-behavioral processes. The recurrence of signatures of selection across Thar populations and their previous identification in other desert-dwelling populations raises the possibility of convergent adaptation. The study also identifies novel candidates for investigating the regulatory and physiological mechanisms underlying adaptation to arid environments. This exploratory study provides a foundation for future studies of interaction between genomic variation, environmental history, and contemporary health in the Thar populations. These findings provide a basis for understanding evolutionary history and exploring the possible implications of adaptive variation under ongoing environmental and lifestyle transitions in Thar communities. Further studies on larger sample sets at genome-wide scales in desert natives and transition populations are required to validate these observations.

## Supporting information

Updated supplemetary figures

## Acknowledgement

We gratefully acknowledge the funding support (I/SEED/MTM/20220130) from IIT Jodhpur and (S/DBT/MTM/PY/20250068) to M.M. from DBT. We extend our thanks to the Department of Bioscience and Bioengineering, IIT Jodhpur, and the HPC facility at IIT Jodhpur. The financial assistance received in the form of student fellowships from the Ministry of Education (MoE), Government of India, to S.M.R. and the University Grants Commission (UGC) to R.J.

## Data Availability

Dataset(s) supporting this study can be found in the Indian Biological Data Centre (IBDC) through the Indian Array Data Archive (IADA) https://ibdc.dbt.gov.in/ under accession number IADA-STmvlrw4hf.

## Authors Contributions

MM conceived the study. SMR performed the experiments and population data analysis with inputs from MM and RJ. SMR, RJ, and MM wrote the manuscript. All authors reviewed and approved the final version of the manuscript.

## Declaration of generative AI and AI-assisted technologies in the writing process

During the preparation of work, we used ChatGPT to improve the text’s readability and language, we reviewed and edited the content as needed, and we took responsibility for it.

