## Supplementary material for "Convergent Genomic Signatures of Selection in Thar: Genetic trade-offs for desert survival": Updated supplemetary figures

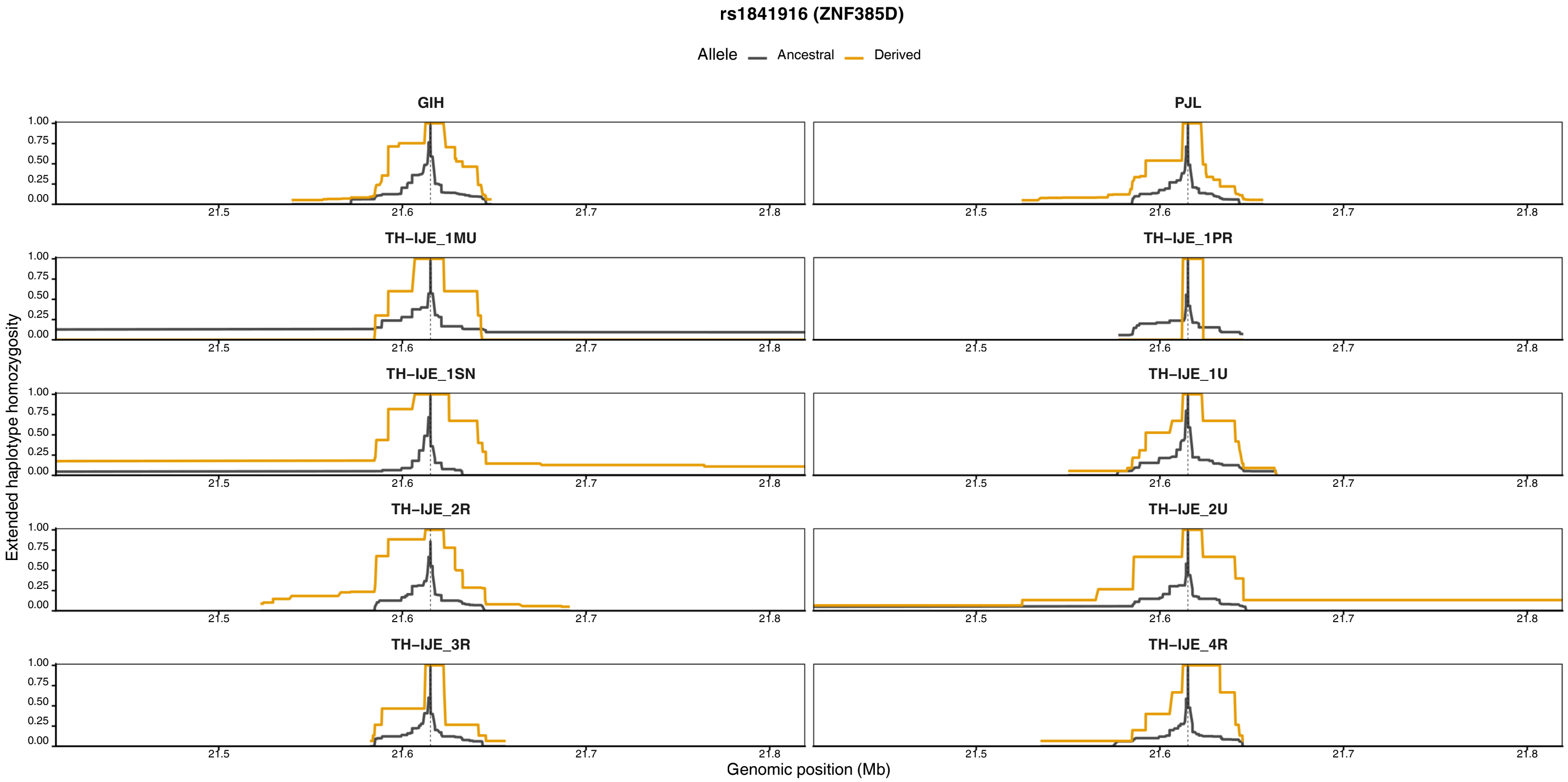
rs1841916 (ZNF385D)
Allele Ancestral Derived
GIH
PJL
TH–IJE_1MU
TH–IJE_1PR
TH–IJE_1SN
TH–IJE_1U
TH–IJE_2R
TH–IJE_2U
TH–IJE_3R
TH–IJE_4R
1.00
0.75
0.50
0.25
0.00
21.5
21.6
21.7
21.8
Extended haplotype homozygosity
Genomic position (Mb)

A)

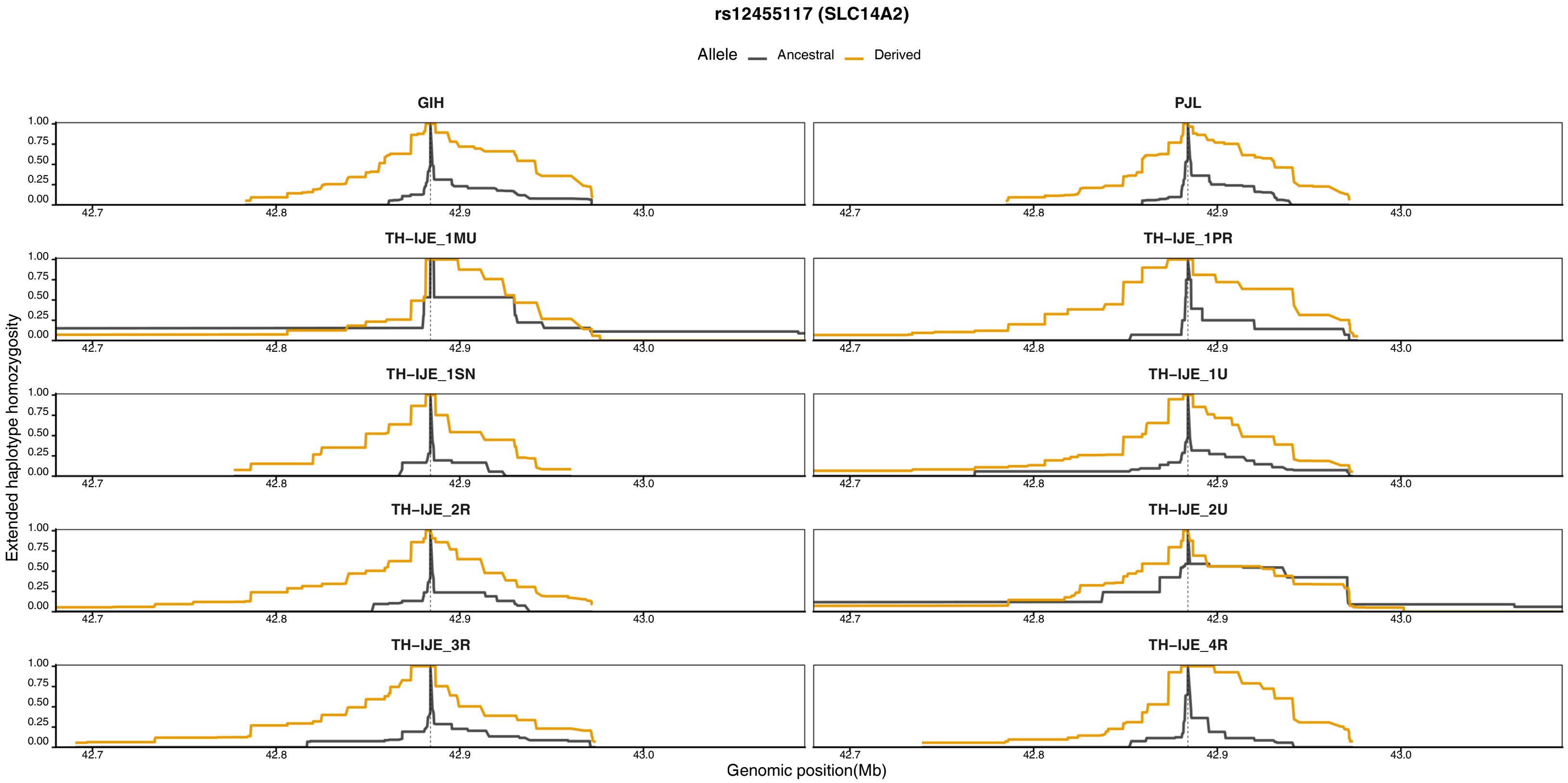
rs12455117 (SLC14A2)
Allele Ancestral Derived
GIH
PJL
TH–IJE_1MU
TH–IJE_1PR
TH–IJE_1SN
TH–IJE_1U
TH–IJE_2R
TH–IJE_2U
TH–IJE_3R
TH–IJE_4R
1.00
0.75
0.50
0.25
0.00
42.7
42.8
42.9
43.0
Extended haplotype homozygosity
Genomic position(Mb)

B)

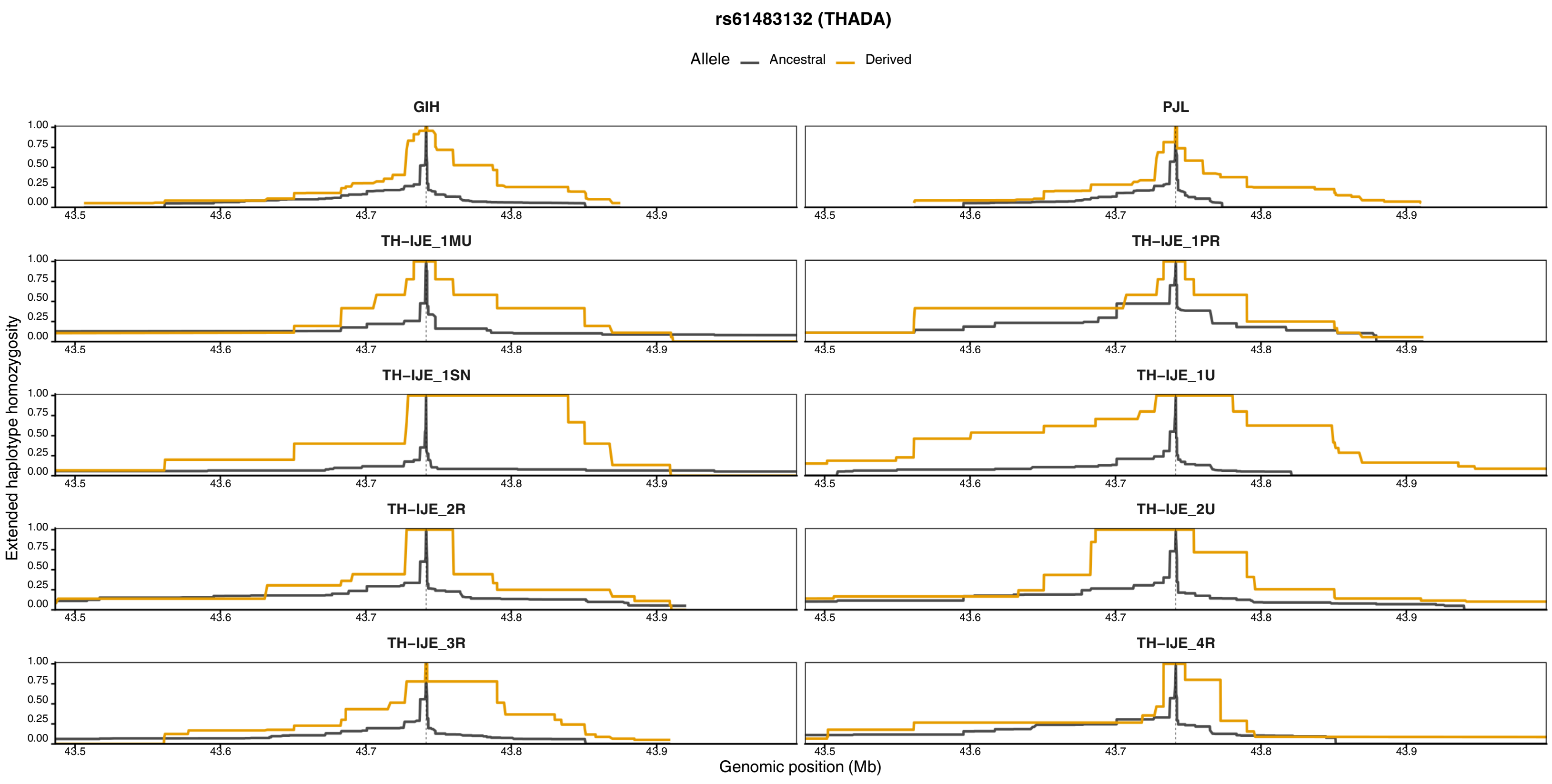
rs61483132 (THADA)
Allele
Ancestral
Derived
GIH
PJL
TH–IJE_1MU
TH–IJE_1PR
TH–IJE_1SN
TH–IJE_1U
TH–IJE_2R
TH–IJE_2U
TH–IJE_3R
TH–IJE_4R
1.00
0.75
0.50
0.25
0.00
43.5
43.6
43.7
43.8
43.9
Extended haplotype homozygosity
Genomic position (Mb)

C)

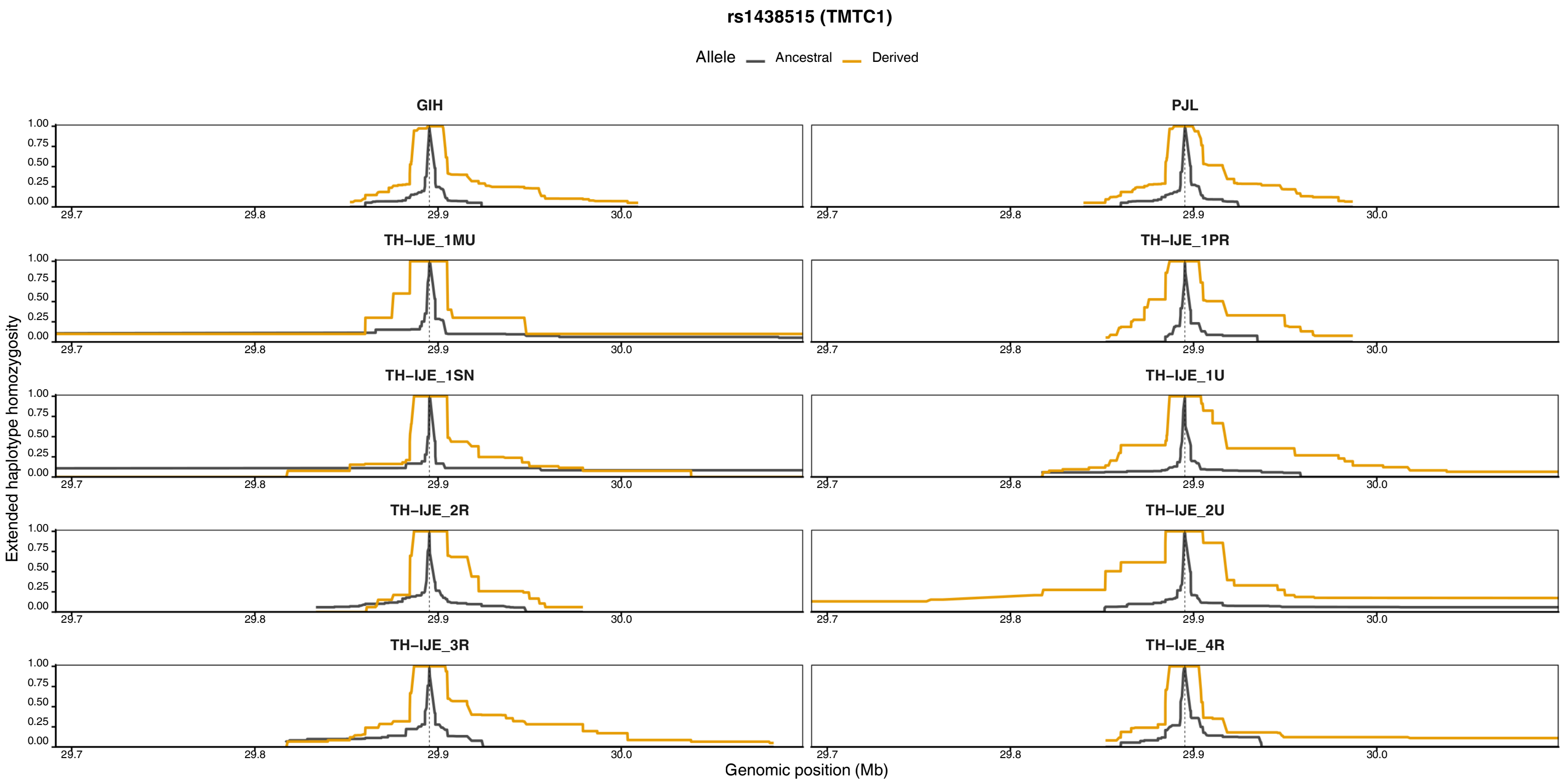
rs1438515 (TMTC1)
Allele
Ancestral
Derived
GIH
PJL
TH–IJE_1MU
TH–IJE_1PR
TH–IJE_1SN
TH–IJE_1U
TH–IJE_2R
TH–IJE_2U
TH–IJE_3R
TH–IJE_4R
1.00
0.75
0.50
0.25
0.00
29.7
29.8
29.9
30.0
Extended haplotype homozygosity
Genomic position (Mb)

D)

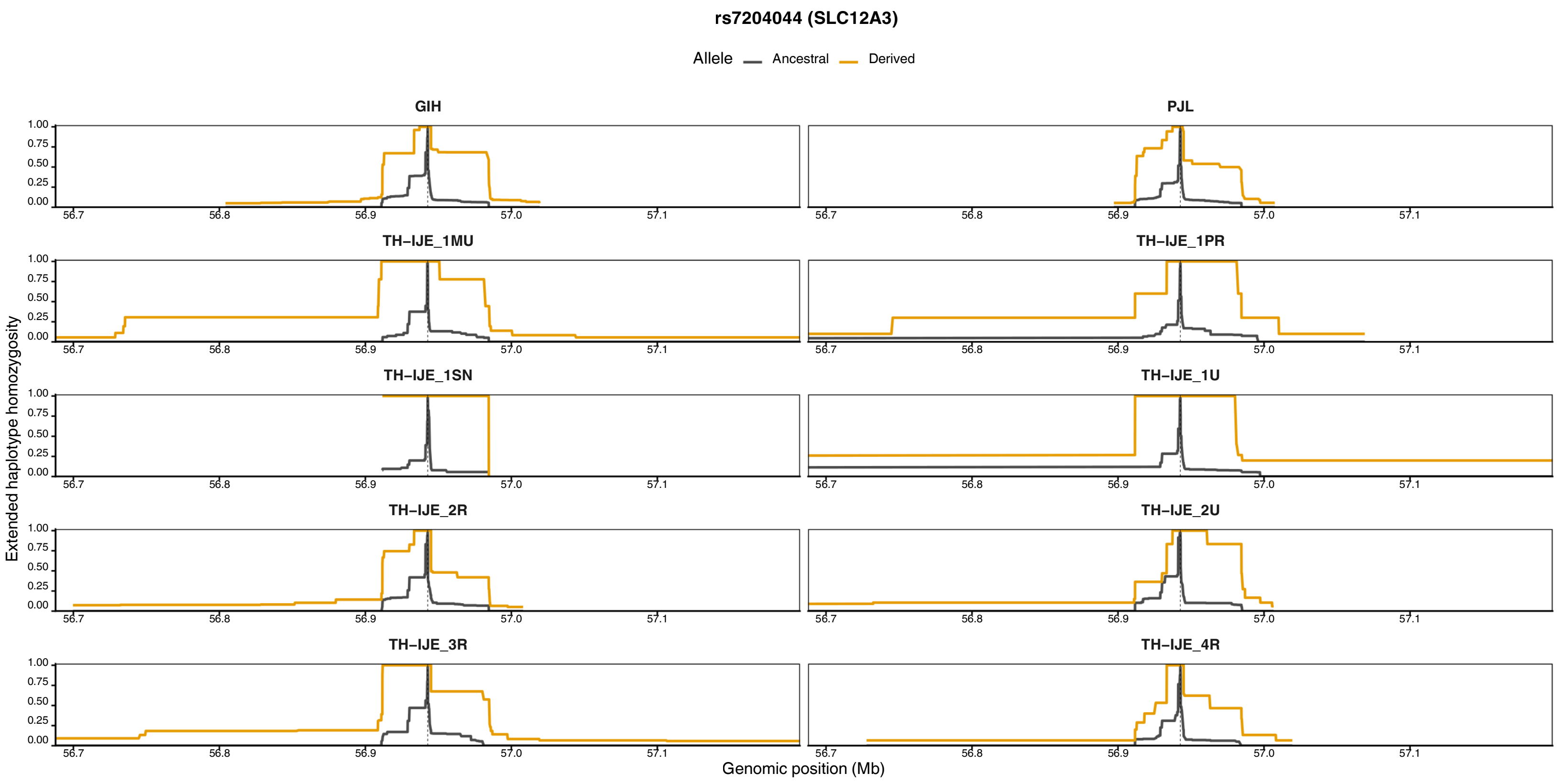
rs7204044 (SLC12A3)
Allele
Ancestral
Derived
GIH
PJL
TH–IJE_1MU
TH–IJE_1PR
TH–IJE_1SN
TH–IJE_1U
TH–IJE_2R
TH–IJE_2U
TH–IJE_3R
TH–IJE_4R
Extended haplotype homozygosity
Genomic position (Mb)
1.00
0.75
0.50
0.25
0.00
56.7
56.8
56.9
57.0
57.1

**E)**

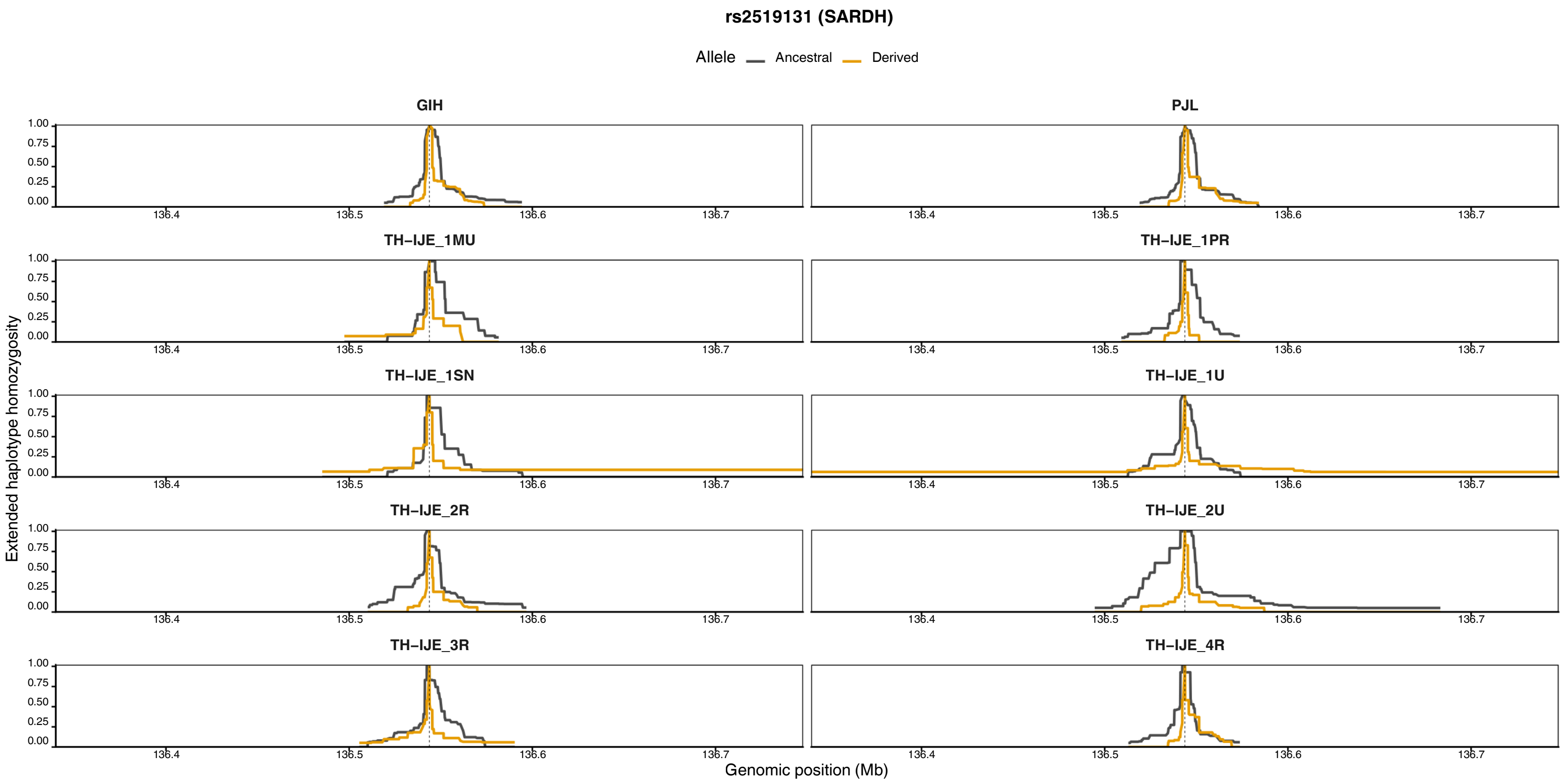
rs2519131 (SARDH)
Allele
Ancestral
Derived
GIH
PJL
TH–IJE_1MU
TH–IJE_1PR
TH–IJE_1SN
TH–IJE_1U
TH–IJE_2R
TH–IJE_2U
TH–IJE_3R
TH–IJE_4R
Extended haplotype homozygosity
Genomic position (Mb)
1.00
0.75
0.50
0.25
0.00
136.4
136.5
136.6
136.7

**F)**

**Supplementary Figure 1. Extended Haplotype Homozygosity (EHH) decay plots for selected candidate SNPs across the Thar populations and reference populations (GIH and PJL).** Extended Haplotype Homozygosity (EHH) decay plots showing the decay of haplotype homozygosity around the focal SNPs in A) ZNF385D (rs1841916), B) SLC14A2 (rs12455117), C) THADA (rs61483132), D) TMTC1 (rs1438515), E) SLC12A3 (rs7204044), and F) SARDH (rs2519131) across the eight Thar populations and the two reference populations (GIH and PJL). The orange line represents the derived allele, while the grey line represents the ancestral allele. The vertical dashed line indicates the focal SNP position.

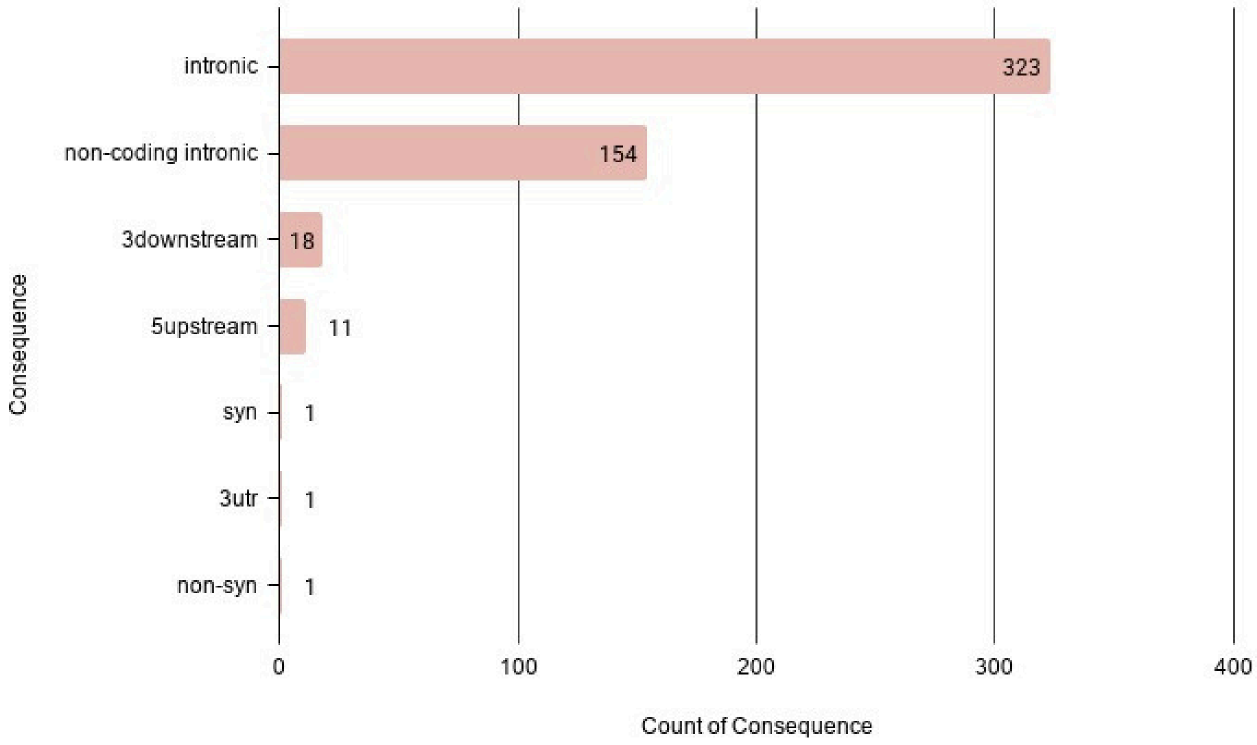


**Supplementary Figure 2. Functional consequence annotation of the 348 top SNPs.** The x-axis represents the count of consequences, and the y-axis represents the variant consequence category.

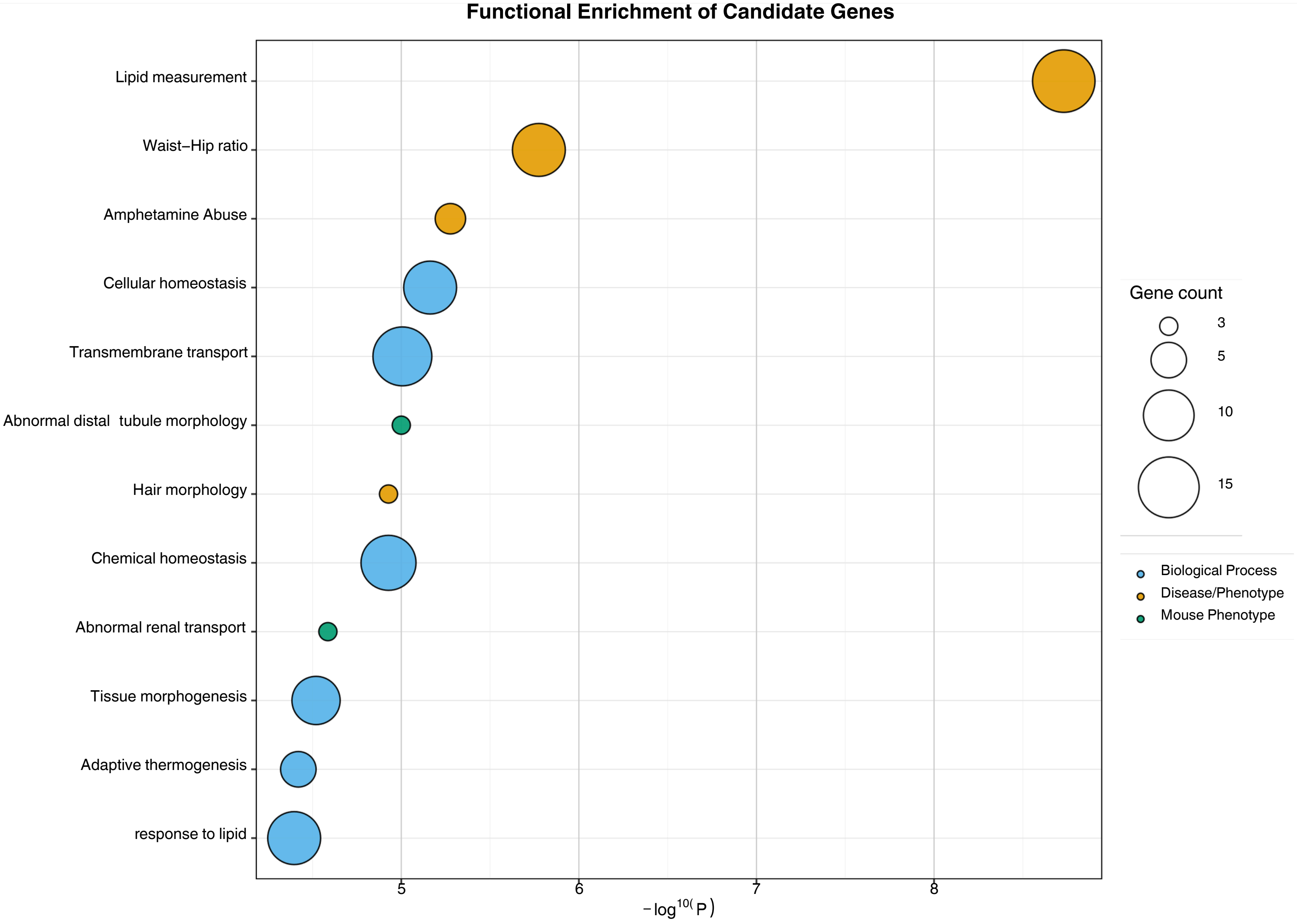


**Supplementary Figure 3. Functional enrichment analysis of top selected candidate genes.** A bubble plot showing the significantly enriched functional categories of candidate genes harboring the 348 putatively adaptive SNPs. The x-axis represents enrichment significance as $-\log_{10}(P)$, while the y-axis shows the enriched terms. Bubble size corresponds to the number of genes associated with each term, and bubble color denotes the enrichment category, including Biological Process, Disease/Phenotype, and Mouse Phenotype.

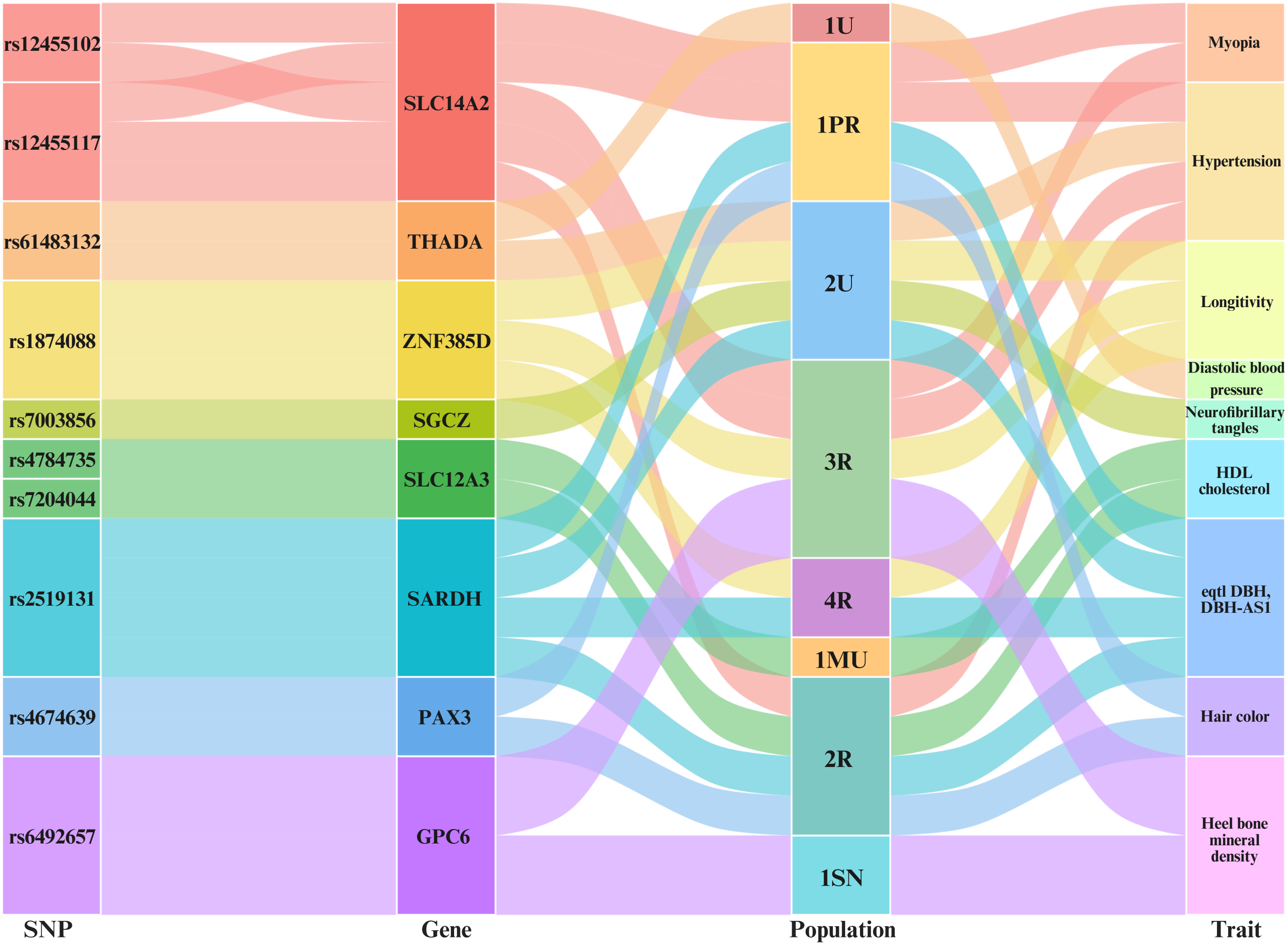


**Supplementary Figure 4.** Diagram of top selection-candidate SNPs, their host genes, the Thar populations in which they showed the strongest selection signal, and their associated phenotypic traits identified through PheWAS, GWAS Catalog, and eQTL analyses.

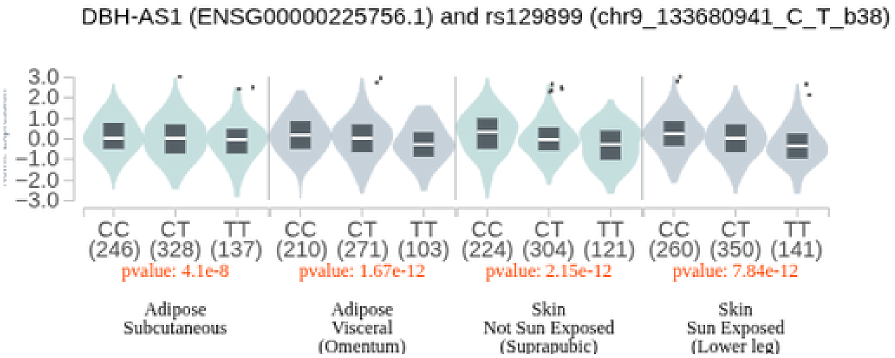


**A)**

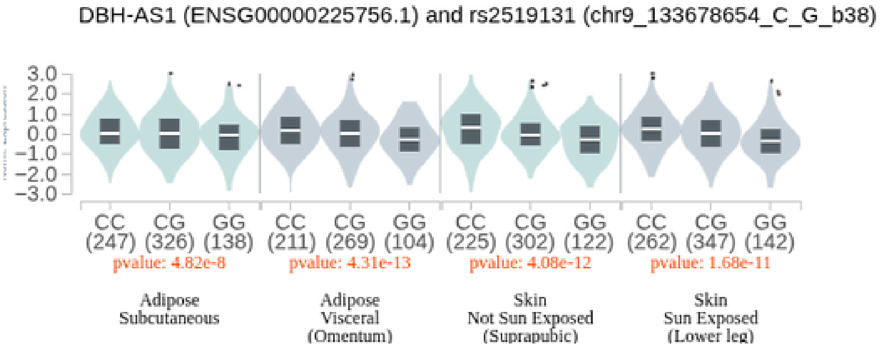


**B)**

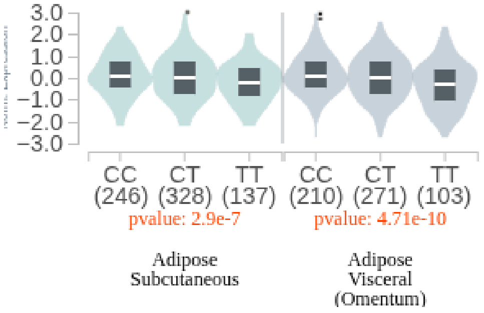


**C)**

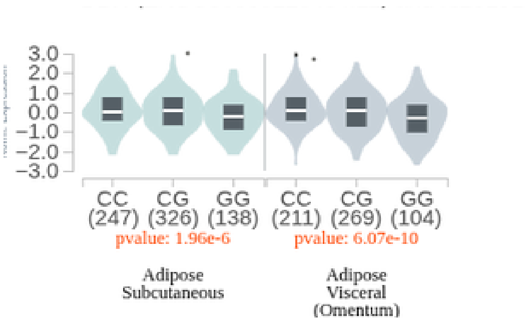


**D)**

**SupplementaryFigure 5.** GTEx cis-eQTL results for the SARDH locus showing the effect of A) rs129899and B) rs2519131 on the expression of the adjacent long non-coding RNA gene DBH-AS1.C) rs129899 and D) rs2519131 on the expression of the adjacent gene DBH.

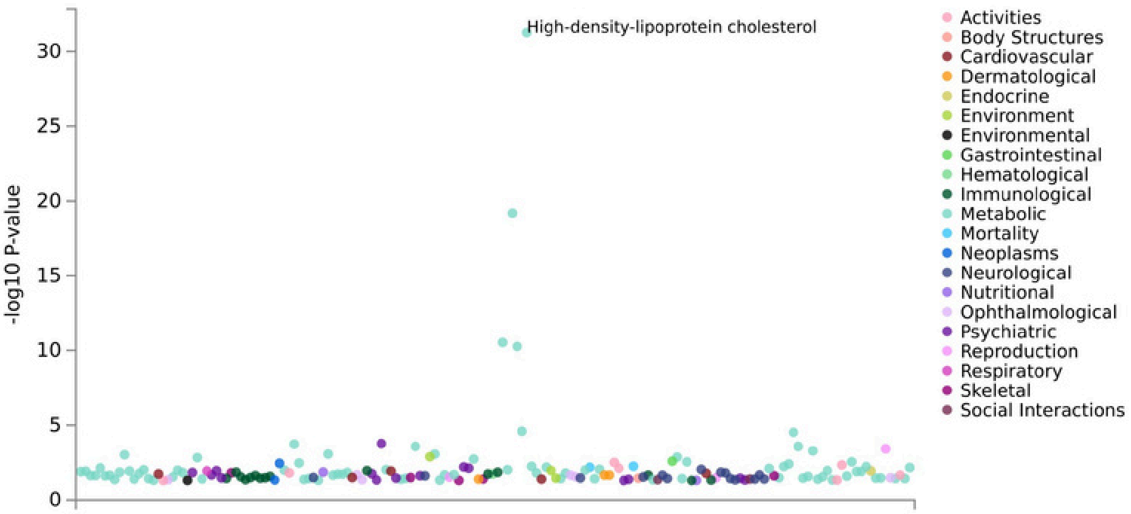


**A)**

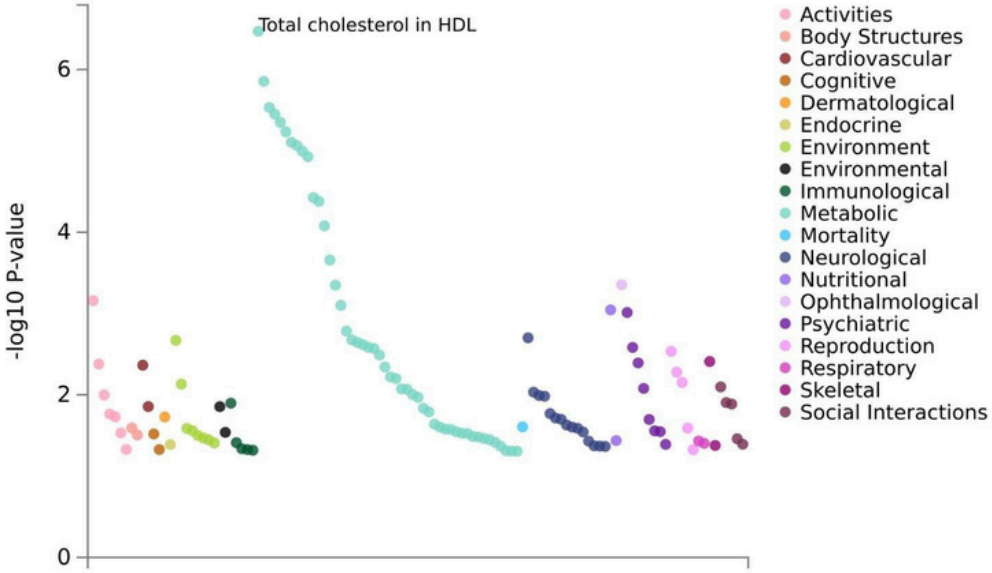


**B)**

**Supplementary Figure 6.** GWAS Atlas phenome-wide association plot for A) rs7204044-[A] and B) rs62035950-[C] of the SLC12A3 locus. Each point represents an association with a phenotypic trait category available in GWASatlas.

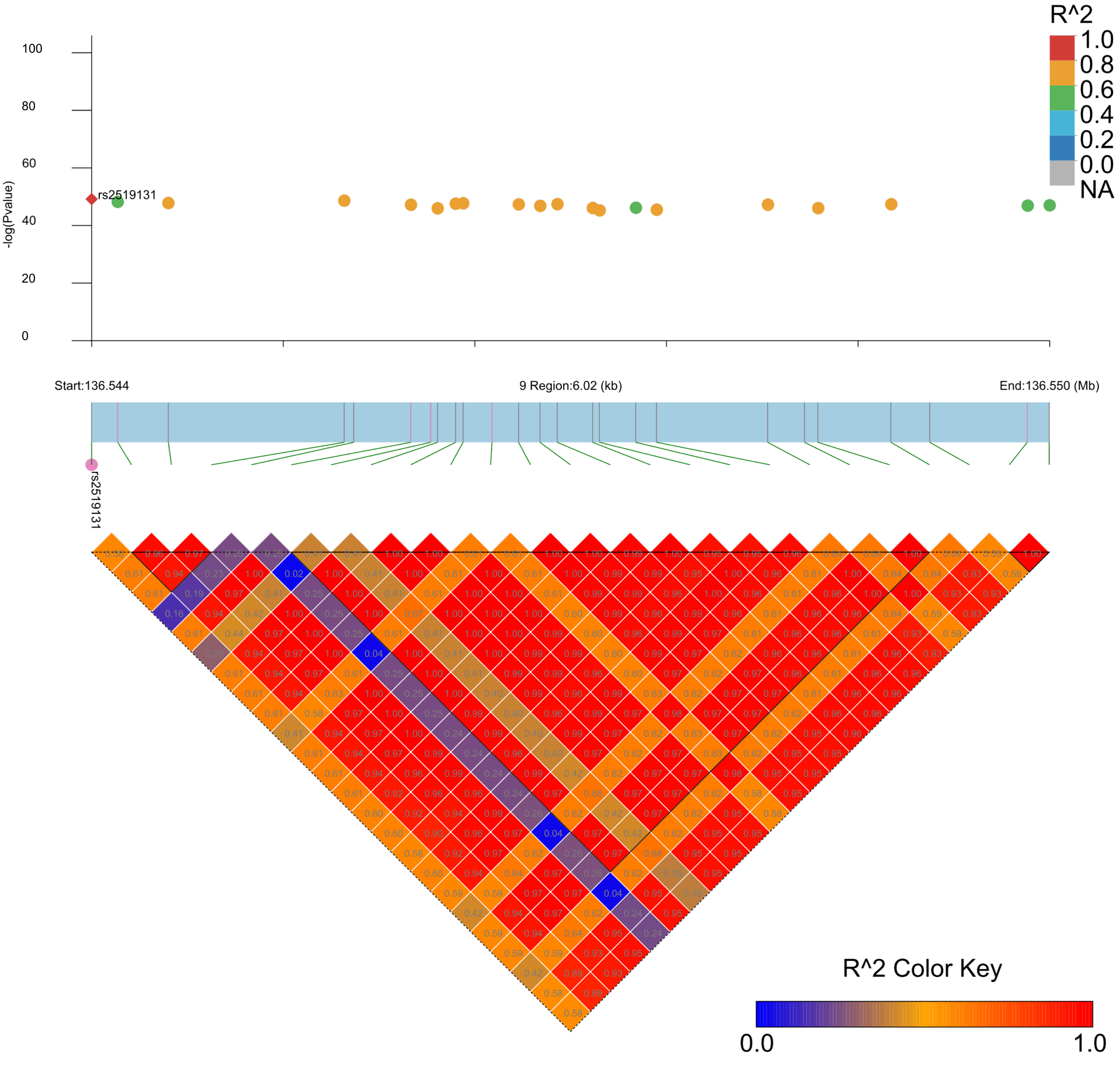
R^2
1.0
0.8
0.6
0.4
0.2
0.0
NA
100
80
60
40
20
0
-log(Pvalue)
rs2519131
Start:136.544
9 Region:6.02 (kb)
End:136.550 (Mb)
rs2519131
R^2 Color Key
0.0
1.0

**A)**

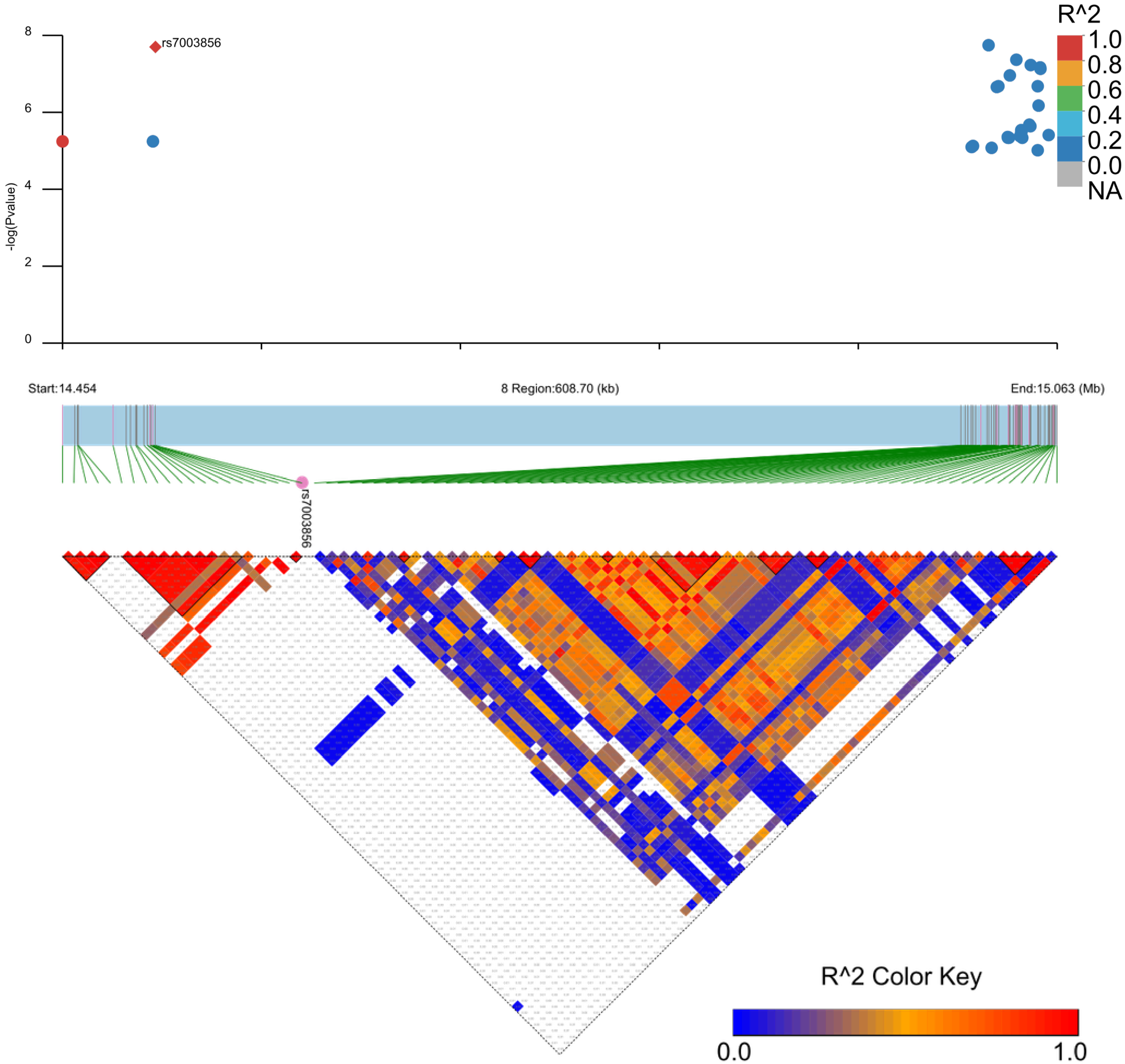
rs7003856
-log(Pvalue)
0
2
4
6
8
R^2
1.0
0.8
0.6
0.4
0.2
0.0
NA
Start:14.454
8 Region:608.70 (kb)
End:15.063 (Mb)
rs7003856
R^2 Color Key
0.0
1.0

**B)**

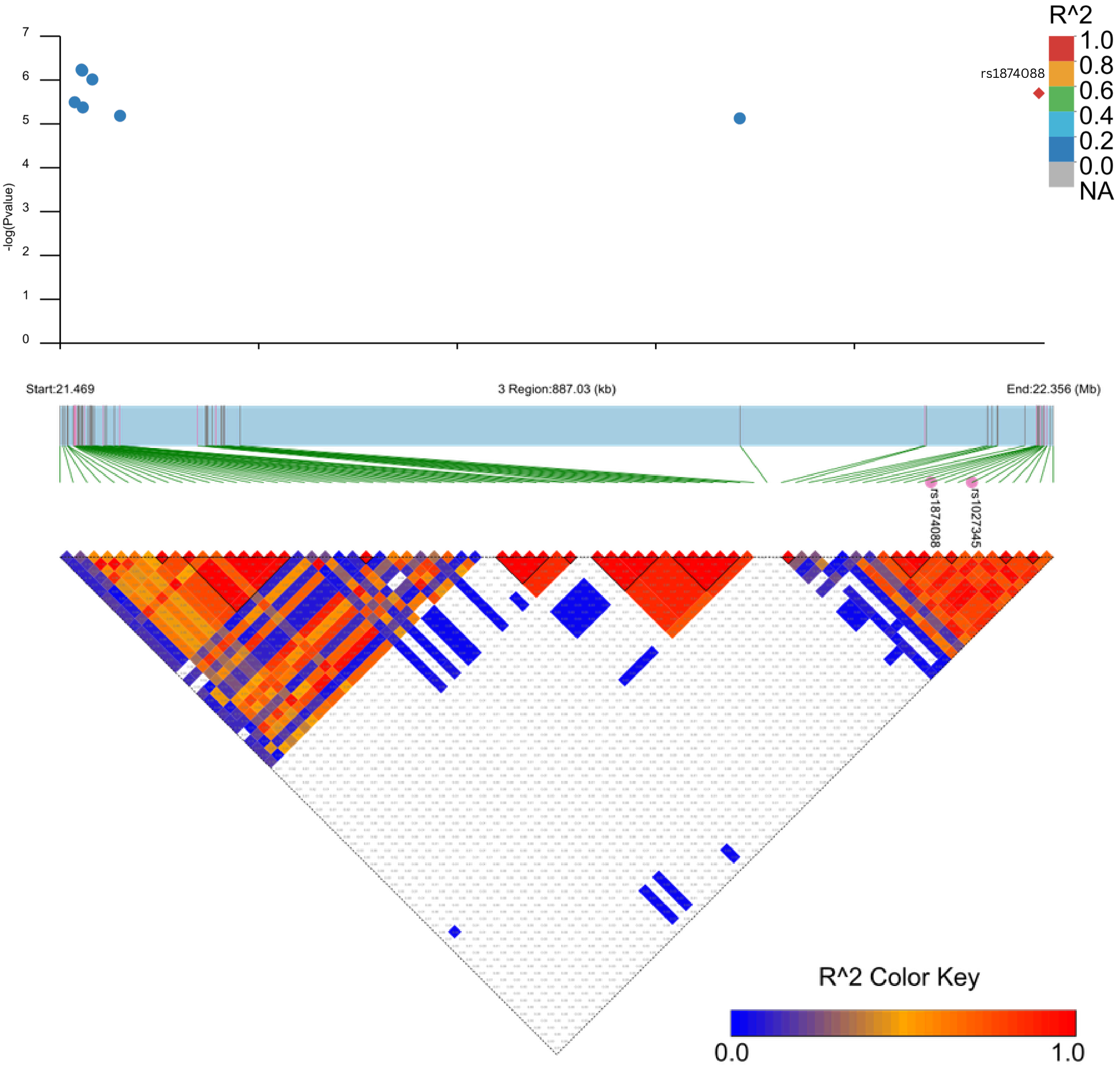


**C)**

**Supplementary Figure 7.** Regional association and LD plots of top selection candidate SNPs: The lower triangle displays pairwise linkage disequilibrium (LD) structure among neighboring variants; the upper panel shows $-\log_{10}$(p-values) of associations retrieved from the multi-database GWAS/PheWAS analysis. A) Regional plot of the SARDH locus encompassing 20 SNPs identified as expression quantitative trait loci (eQTLs), with rs2519131 highlighted as the top selection candidate. B) Regional plot of SGCZ highlighting rs7003856 associated with neurofibrillary tangles. C) Regional plot of the ZNF385D locus showing rs1874088 associated with longevity.

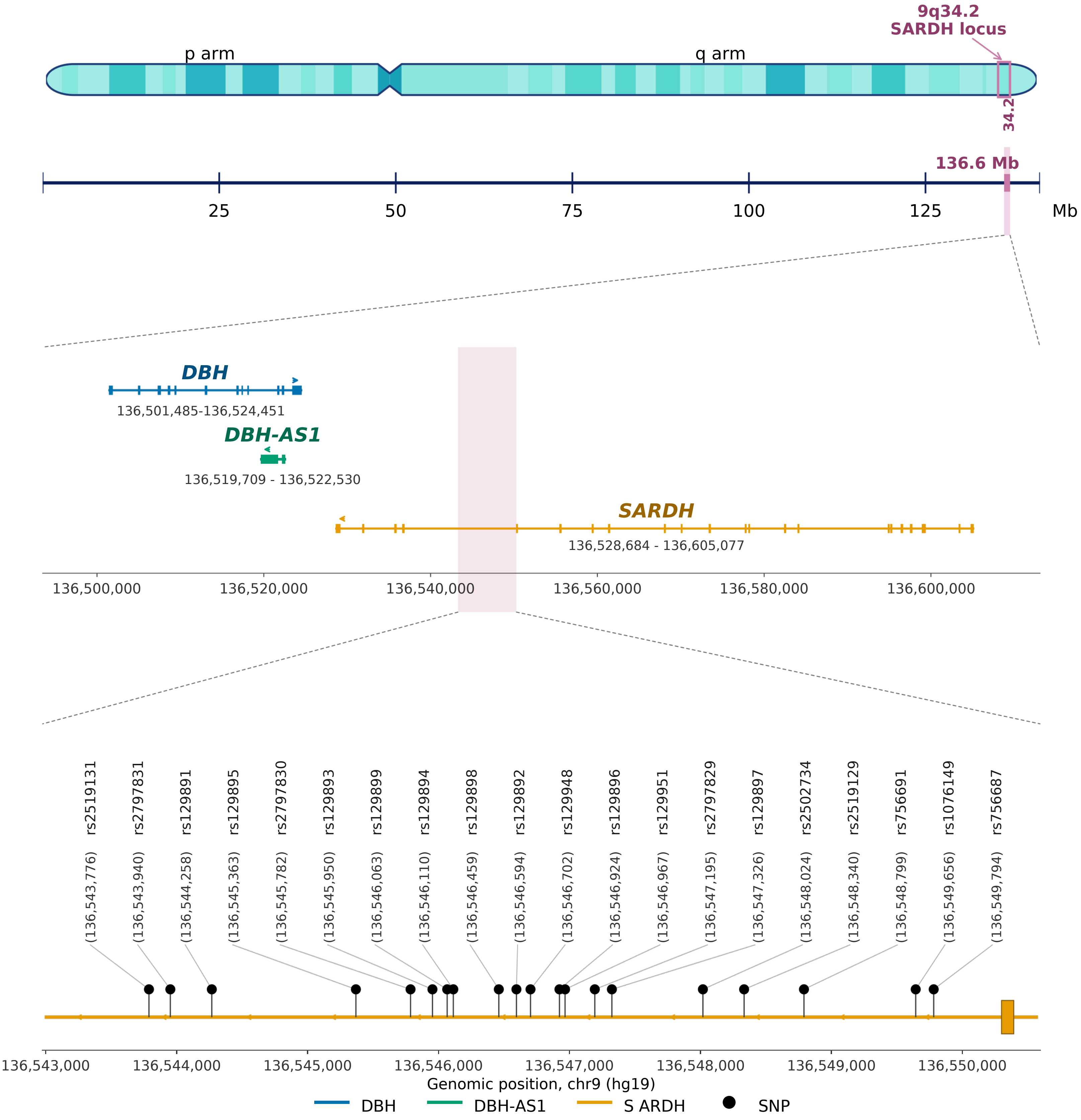


**Supplementary Figure 8.** Location of the top 20 SARDH SNPs on chromosome 9q34.2 (hg19) relative to the neighboring DBH and DBH-AS1 genes. Dashed lines indicate the zoomed region.
